# *Legionella pneumophila*’s Tsp is important for surviving thermal stress in water and inside amoeba

**DOI:** 10.1101/2020.12.08.417196

**Authors:** Joseph Saoud, Thangadurai Mani, Sébastien P. Faucher

## Abstract

*Legionella pneumophila* (*Lp*) is an inhabitant of natural and man-made water systems where it replicates within amoebae and ciliates and survives within biofilms. When *Lp*-contaminated aerosols are breathed in, *Lp* will enter the lungs and infect human alveolar macrophages, causing a severe pneumonia known as Legionnaires Disease. *Lp* is often found in hot water distribution systems (HWDS), which are linked to nosocomial outbreaks. Heat treatment is used to disinfect HWDS and reduce the concentration of *Lp*. However, *Lp* is often able to recolonize these water systems, indicating an efficient heat-shock response. Tail-specific proteases (Tsp) are typically periplasmic proteases implicated in degrading aberrant proteins in the periplasm and important for surviving thermal stress. In this paper, we show that Tsp, encoded by the *lpg0499* gene in *Lp* Philadelphia-1, is important for surviving thermal stress in water and for optimal infection of amoeba when a shift in temperature occurs during intracellular growth. Tsp is expressed in the post-exponential phase but repressed in the exponential phase. The *cis*-encoded small regulatory RNA Lpr17 shows opposite expression, suggesting that it represses translation of *tsp*. In addition, *tsp* is regulated by CpxR, a major regulator in *Lp*, in a Lpr17-independent manner. Deletion of CpxR also reduced the ability of *Lp* to survive heat shock. In conclusion, this study shows that Tsp is an important factor for the survival and growth of *Lp* in water systems.

**IMPORTANCE:** *Legionella pneumophila* (*Lp*) is a major cause of nosocomial and community-acquired pneumonia. *Lp* is found in water systems including hot water distribution systems. Heat treatment is a method of disinfection often used to limit *Lp*’s presence in such systems; however, the benefit is usually short term as *Lp* is able to quickly recolonize these systems. Presumably, *Lp* respond efficiently to thermal stress, but so far not much is known about the genes involved. In this paper, we show that the Tail-specific protease (Tsp) and the two-component system CpxRA are required for resistance to thermal stress, when *Lp* is free in water and when it is inside host cells. Our study identifies critical systems for the survival of *Lp* in its natural environment under thermal stress.

## INTRODUCTION

Legionnaires’ disease (LD) is a severe form of pneumonia in human caused by the Gram-negative bacterium *Legionella pneumophila* (*Lp*) (1). *Lp* is often responsible for nosocomial and community-acquired pneumonia (1). In the environment, *Lp* can be found in natural and man-made aquatic environments, such as cooling towers and water distribution systems, where it can replicate within phagocytic protozoa (2–4). Notably, *Lp* replicates within *Vermamoeba vermiformis* (formerly *Hartmannella vermiformis*), a thermotolerant amoeba commonly found in both natural and man-made water systems (5, 6). *V. vermiformis* protects *Lp* from predation, competition, and various disinfection methods such as heat treatments, potentially contributing to nosocomial infections (3, 5, 6). In addition, intracellular growth of *Lp* within amoeba increases its pathogenicity and facilitates the establishment of an infection in humans (3). The main virulence factor of *Lp* is the Type IVb Secretion System called Icm/Dot, which translocate more than 300 effectors inside the host cells (7). These effectors are responsible for stopping the maturation of the phagosome and creating a specialized vacuole where *Lp* can grow (7).

The preferred growth temperature of *Lp* is between 25 °C and 42 °C, though it has been found in water systems at temperatures below 20 °C or above 60 °C (8). Its ability to live in a wide range of temperature allows *Lp* to colonize hot water distribution systems (HWDS) (9). The higher temperature found within HWDS also reduces the microbial diversity, therefore making it easier for *Lp* to compete in these environments (10, 11). A method used to limit the proliferation of *Lp* in water systems is to maintain the temperature at the outlet above 55 °C (8, 9). When a system is heavily colonized by *Lp*, it can be treated by superheat and flush, also referred to as pasteurization, consisting of increasing the temperature at the heater to about 75 °C in order to provide a temperature of at least 65 °C at the outlets (12).

An increase in temperature, even by a few degrees, can cause proteins to unfold which leads to protein aggregates that can be lethal as these aggregates accumulate in the cells (13, 14). Protein misfolding and aggregation results in an imbalance of protein homeostasis, which triggers the heat shock response (14). The canonical heat shock response is composed of two heat shock regulons, the σ^H^ and σ^E^ heat shock regulons (14). This response results in the production of chaperones and proteases to refold or destroy misfolded proteins and aggregates (14). In *Lp*, the two-component system LetA/S is the only regulatory system known to be important for surviving thermal stress (15).

Protein degradation by proteases is an important cellular function as it allows the removal of aberrant proteins, the regulation of intracellular protein concentration, produce active molecules from precursors such as the a-toxin from *Staphylococcus aureus* or the heat-stable toxin produced by enterotoxigenic *Escherichia coli* (ETEC), and allows the cell to recycle amino acids during starvation (16, 17). Carboxyl-terminal proteases (CTPs) are serine proteases conserved in most Gram-negative bacteria (18–20). They can also be found in archaea, Gram-positive bacteria, eukaryotes, viruses, as well as in organelles such as chloroplasts (20–22). CTPs are mostly located in the periplasm though some are located in the cytoplasm, while others are secreted in the extracellular environment (19). CTP proteases are involved in several different processes in Gram-negative bacteria. *E. coli*’s CTP, called tail-specific protease (Tsp) and sometimes Prc, was the first bacterial CTP characterized and is involved in regulating peptidoglycan assembly (19). The assembly of the peptidoglycan layer play a role in the bacteria’s size and shape (23). This important process is dependent on penicillin-binding proteins (PBP) (20, 24). Tsp recognizes a sequence of amino acids located on the C-terminal end of the precursor of PBP-3, resulting in its activation (25, 26). As a result, the *E. coli*’s *tsp* mutant is sensitive to various antibiotics, thermal and osmotic stress, and displays a filamentous morphology (25, 27). Overproduction of Tsp is detrimental to cell growth (25). The susceptibility to thermal stress is dependent on the osmolarity. These phenotypes are attributed to an increased permeability of the outer membrane in the *tsp* mutant (25, 28). The *tsp* mutant and the WT expressed similar levels of GroEL and DnaK, two heat shock proteins, when exposed to thermal stress in high osmolarity buffer (25). However, when the strains were in low osmolarity buffer and exposed to thermal stress, the *tsp* mutant showed decreased expression of DnaK and almost no expression of GroEL (25). *E. coli*’s Tsp also targets MepS, a peptidoglycan hydrolase which cleaves peptide cross-links between the glycan chains of the peptidoglycan layer (29, 30). Tsp is also able to degrade misfolded proteins in the periplasm, suggesting a role in maintaining protein homeostasis in the periplasm (26, 31, 32).

*Pseudomonas aeruginosa* codes for two CTPs, called Prc and CtpA (19, 33). Prc is a homologue of *E. coli*’s Tsp and has been shown to degrade a mutant form of the anti-sigma factor MucA, preventing development of mucoidy (33–35). CtpA is involved in the proper function of the type 3 secretion system (T3SS), for cytotoxicity in cultured cells and for virulence in the animal model of acute pneumonia (19, 33). The activity of CtpA is dependent on LcbA, an outer membrane lipoprotein with tetratricopeptide repeat (TPR) motifs (36). CtpA degrades four peptidoglycan hydrolases: MepM and PA4404, belonging to the M23 peptidase family, and PA1198 and PA1199, belonging to the NlpC/P60 peptidase family (36).

The *lpg0499* gene in *Lp* encodes a CTP protease named Tsp, most homologous to the *P. aeruginosa* CtpA (37). The *tsp* gene is upregulated 54-fold in the post-exponential phase compared to the exponential phase (38). The *tsp* gene was also strongly upregulated during infection of *Acanthamoeba castellanii* and THP-1 (38, 39). *tsp* is regulated by the *Legionella* quorum sensing system, an important activator of genes required in the transmissive phase (40). A transcriptomic analysis of a *letS* mutant in water revealed that *tsp* is downregulated in the mutant (15). In addition, the expression of *tsp* was downregulated in a *cpxR* mutant grown to PE phase, suggesting CpxR is also required for expression of *tsp* in PE phase (41). CpxR regulates the expression of several Icm/Dot effectors and is required for growth in *A. castellanii* (41–44). A previous study showed that a *tsp* mutant in the *Lp* serogroup 1 strain 130b did not have a growth defect in *Acanthamoeba castellanii*, in THP-1 macrophages or in low-salt chemically defined media at 42°C (37). A small regulatory RNA (sRNA) named Lpr17 (Lppnc0140 in strain Paris) is encoded complementary to *lpg0499* (45, 46). sRNA are short RNA molecules involved in post-transcriptional regulation of genes required for virulence and response to various conditions such as sugar metabolism, iron homeostasis, and biofilm formation (47, 48). Base-pairing sRNAs are the most common type of sRNA and they act by hybridising to their target mRNA. Lpr17 is a *cis*-encoded sRNA. Such sRNA pair perfectly with their target due to them being encoded on the complementary strand of their target (47). Lpr17 is therefore likely to control *tsp* expression.

In this paper, we have investigated the role of *Lp*’s Tsp in the resistance to thermal stress in water and during infection of amoeba. In addition, we have studied the regulatory function of the *cis*-encoded sRNA Lpr17 as well as confirmed the role of CpxR in the regulation of Tsp.

## RESULTS

### Tsp is important for *L. pneumophila* to survive thermal stress

Since Tail-specific proteases have been implicated in managing thermal stress in other bacteria, the importance of Tsp for the survival of *Lp* after a heat shock at 55 °C was tested (Figure 1). The strains were suspended in Fraquil, an artificial freshwater medium, and incubated for 24 hours at room temperature prior to the temperature stress. After 15 minutes at 55 °C, the CFU count of the Δ*tsp* mutant decreased by 10,000-fold while the CFU count of the WT and the complemented strain decreases by only 10-fold. 30 minutes after thermal stress, the CFU count of the Δ*tsp* mutant was 100 times lower than the CFU count of the WT and complemented strain. However, the difference between the mutant and the WT was considered statistically significant only at 15 minutes.

**Figure 1:**
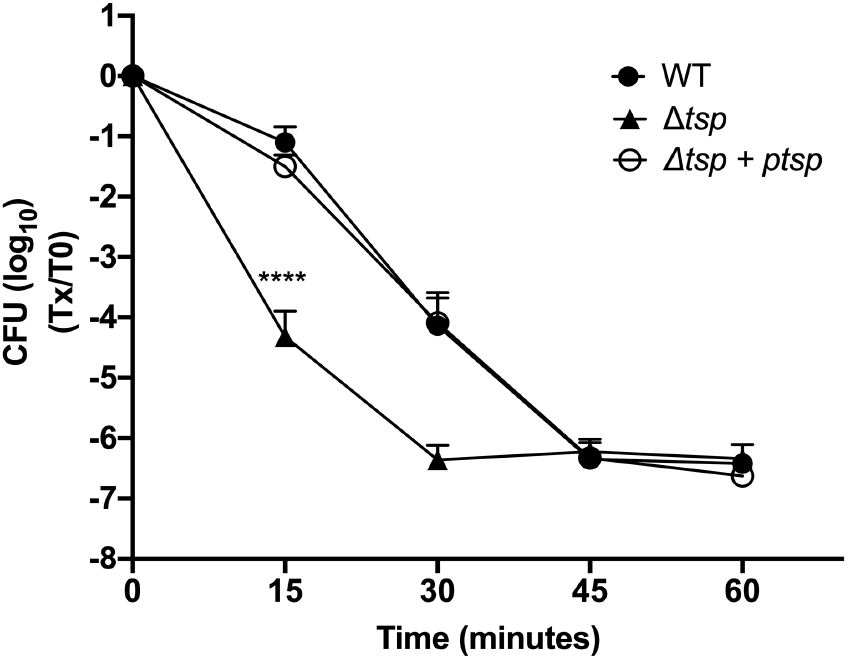
Tsp is required for *Lp* to survive a thermal stress in water. The WT, *tsp* mutant (Δ*tsp*), and the complemented strain (Δ*tsp* + p*tsp*) were suspended in Fraquil for 24 hours and then subjected to thermal stress at 55 °C. The survival of the strains was measured by CFU counts every 15 minutes from 0 to 60 minutes. The data shown represent the average of three independent biological replicates with standard deviation. A two-way ANOVA with a Tukey correction for multiple comparison was used to determine statistical difference (****: *P*-value < 0.0001).

### Tsp is important for intracellular multiplication in *V. vermiformis* following a temperature shift

Given the inability of the Δ*tsp* mutant to survive a heat treatment, we hypothesized that the ability of the mutant to grow inside amoeba could be compromised if a temperature shift occurs, which is likely to happen in a HWDS. The ability of the Δ*tsp* mutant to replicate within *V. vermiformis* was then investigated (Figure 2). The infection was therefore carried out at 3 different temperatures in parallel: room temperature for 5 days, 37 °C for 5 days, and room temperature for 2 days followed by 37 °C for 3 days (referred to in this manuscript as a temperature shift). The temperature shift would simulate the change in temperature encountered in HWDS, albeit to a lower degree, and would test the importance of Tsp for survival under these conditions. At room temperature, none of the strains tested were able to replicate intracellularly (Figure 2A). At 37 °C, all strains were able to replicate intracellularly, with the exception of *dotA^−^*, the negative control (Figure 2C), confirming what was previously reported (37). However, when the temperature was shifted from 25 °C to 37 °C two days after the start of the infection, the Δ*tsp* mutant had a severe growth defect within amoeba compared to the WT (Figure 2B). At the end of the experiment, the mutant showed significantly less intracellular growth than the WT and complemented strain when a temperature shift occurs.

**Figure 2:**
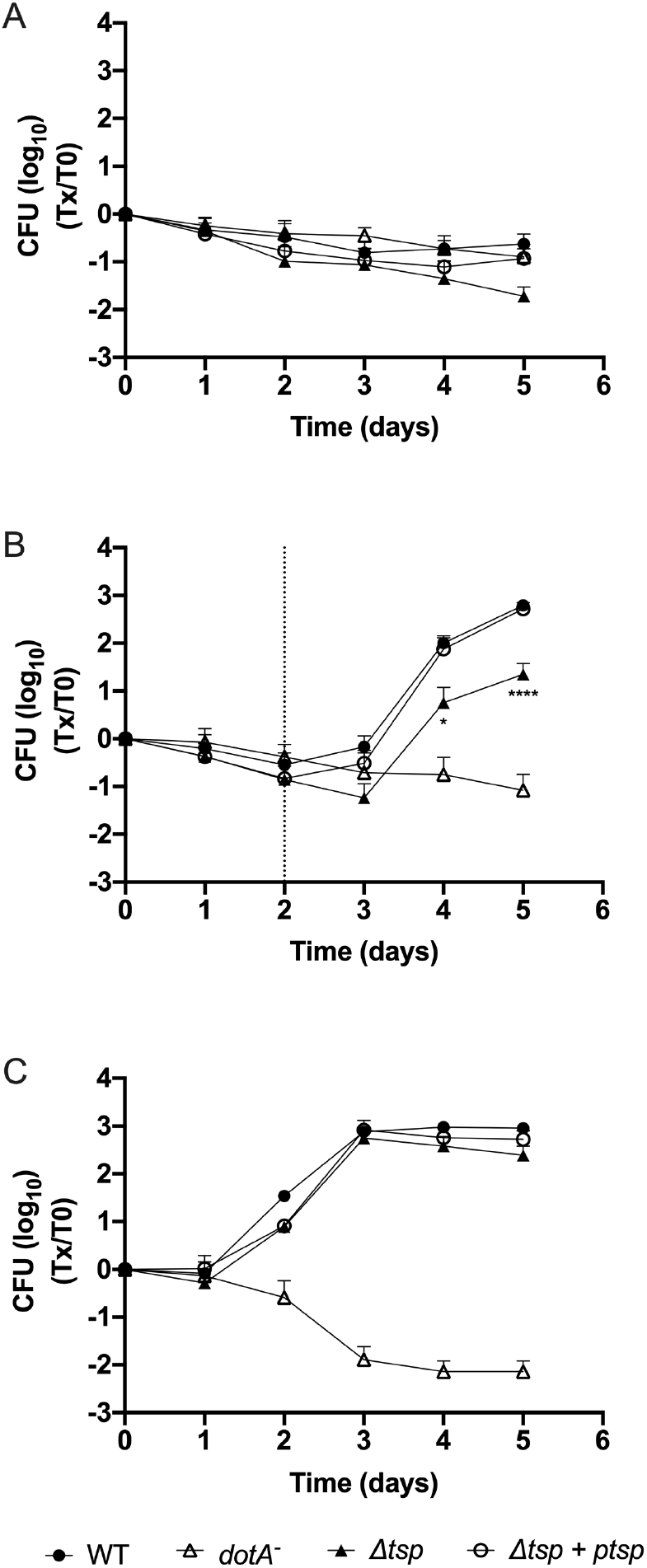
Tsp is required for intracellular multiplication in *V. vermiformis* during a temperature shift. Infection of the host cell *V. vermiformis* was carried out at room temperature for 5 days (A), at room temperature for 2 days then at 37 °C for 3 days (B), and at 37 °C for 5 days (C). Host cells were infected with the WT, *tsp* mutant (Δ*tsp*), complemented strain (Δ*tsp* + p*tsp*), and *dotA^−^*, a negative control. The dotted line in panel B represents the day on which the shift in temperature occurred. The data represent average and standard deviation of 6 biological replicates. A two-way ANOVA with a Tukey correction for multiple comparison was used to determine if the results were statistically different. The statistical significance shown in Figure 2 compares the WT and *tsp* mutant (*, *P*-value < 0.05; ****, *P*-value < 0.0001).

### Lpr17 is expressed in E phase

Next, we sought to investigate the regulation of Tsp. The small regulatory RNA Lpr17 is encoded antisense to the 5’ end and promoter region of the *tsp* gene and the 3’ end of *lpg0500*, which codes for a peptidase of the M23/M37 family (Figure 3A). *Cis*-encoded sRNAs tend to regulate the gene encoded on the complementary stand (47). Therefore, the expression of the Lpr17 sRNA was analyzed by northern blot in the WT, *tsp* mutant (Δ*tsp*), and complemented strain (Δ*tsp* + p*tsp*) (Figure 3B). To make the mutant, the *tsp* gene was replaced with a kanamycin resistance cassette, and, in the process, a section of Lpr17 coding region including its transcription start site (TSS) was also removed. The complementation in *trans* was done by cloning the full-length *tsp* with 441 bp upstream of the ATG, to include the putative promoter on a plasmid. This fragment also contains the full length *lpr17* gene. Lpr17 was only detected in E phase by northern blot. The expression of Lpr17 is stronger in the complemented strain, probably caused by the copy number of the plasmid used for complementation. In the complemented strain, Lpr17 was also expressed in PE phase, although much less than in E phase.

**Figure 3:**
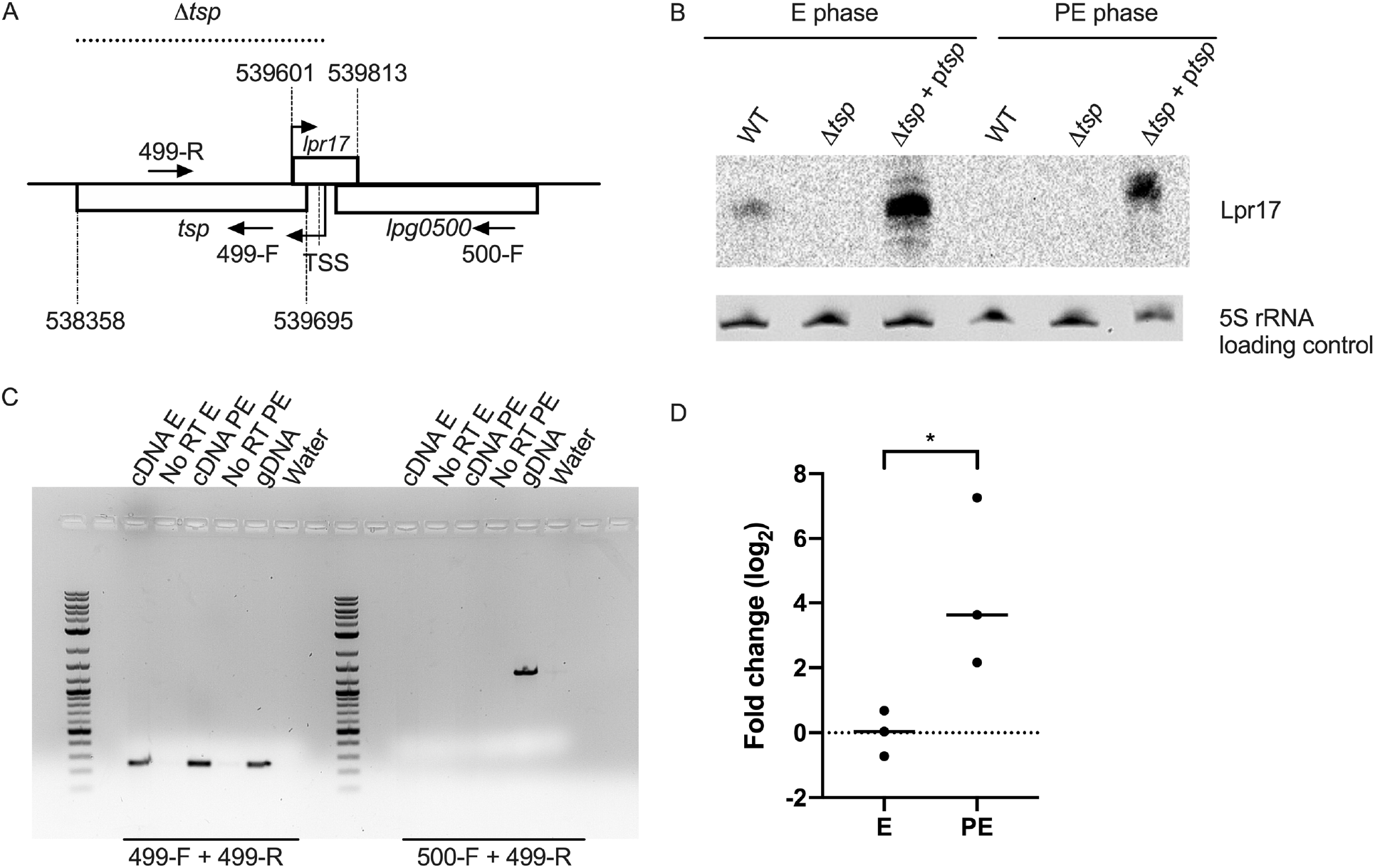
The *cis*-encoded Lpr17 sRNA is expressed in E phase while *tsp* is expressed in both E and PE phase. A) *tsp* is encoded downstream of *lpg0500* and the Lpr17 sRNA is encoded on the complementary strand of *tsp* overlapping with its TSS and the promoter region. The coordinates of *tsp* and *lpr17* in the Philadelphia-1 genome are indicated. The dotted line indicates the portion of the genome that was replaced with a kanamycin resistance cassette in the Δ*tsp* mutant. B) Northern blotting was used to investigate the expression of Lpr17 in the WT, *tsp* mutant (Δ*tsp*), and complemented strain (Δ*tsp* + p*tsp*) grown to exponential phase (E) and post-exponential phase (PE) in AYE broth. 5S rRNA was used as a loading control. C) An RT-PCR was performed on cDNA from the WT strain grown to E and PE phase in AYE broth to determine if *tsp* is encoded on a polycistronic operon with the upstream gene *lpg0500*. RNA from E and PE phase that was not reverse transcribed (No RT) as well as water were used as a negative control, genomic DNA (gDNA) of the WT strain served as a positive control. D) RT-qPCR was used to investigate the expression of *tsp* in the WT grown in AYE broth to PE phase compared to E phase. An unpaired t-test was used to access statistical significance (*, *P* <0.05).

### *tsp* is transcribed in E and PE phase

RT-PCR was used to investigate when the *tsp* gene is transcribed and if the *tsp* and *lpg0500* genes are transcribed as part of a polycistronic mRNA (Figure 3C). Amplification with a forward primer (499-R) and a reverse primer (499-F) within the *tsp* gene revealed that the gene is transcribed in both E and PE phase. The 499-R primer along with a forward primer within *lpg0500* (500-F) showed that the 2 genes are not expressed as a polycistronic mRNA. Our RT-PCR results suggest that *tsp* is slightly more expressed in PE phase than E phase. Using RT-qPCR, *tsp* is 4.3-fold more expressed in PE phase than E phase (Figure 3D).

### Tsp is expressed in PE phase

In order to detect the expression of Tsp, the protein was tagged with a hexahistidine tag and cloned onto pXDC39, a derivative of pMMB207c lacking the *Ptac* promoter and *lacI*^q^ (49). This plasmid also encodes for a full-length copy of *lpr17*. The expression of Tsp and Lpr17 was investigated by western blot and northern blot, respectively, in the WT and the Δ*tsp* mutant harbouring the Tsp-his construct (p*tsp*-his) in E and PE phase (Figure 4). The WT and the Δ*tsp* mutant containing an empty vector served as controls. As expected, the Lpr17 sRNA is more expressed in E phase than PE phase (Figure 4A), similarly to what was seen in the complemented strain (Figure 3B). In contrast, the Tsp protein is only detected in PE phase (Figure 4A).

**Figure 4:**
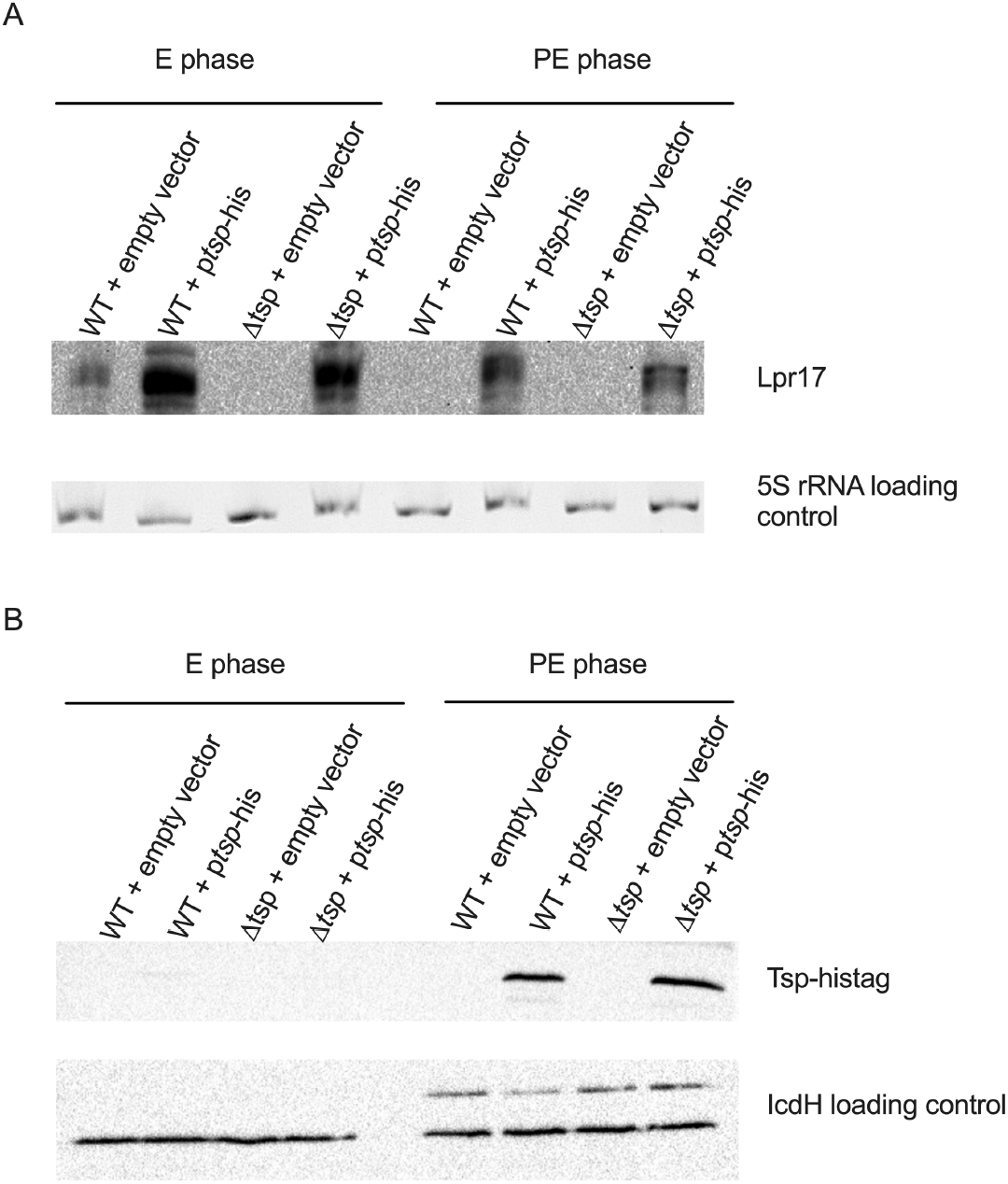
Tsp is expressed in PE phase. Tsp was tagged with a polyhistidine tag and cloned in plasmid pXDC39 under its own promoter and transferred to the WT (WT + p*tsp-his*) and *tsp* mutant (Δ*tsp* + p*tsp-his*) strains. The strains were grown to E and PE phase in AYE broth, and expression of the Lpr17 was analyzed by northern blot (A) while the expression of the Tsp was analyzed by western blot (B). The strains with the empty vector (WT + empty vector; Δ*tsp* + empty vector) served as negative controls. 5S rRNA and IcdH serve as the loading control for northern blot and western blot, respectively.

### Lpr17 is repressed during thermal stress

Lpr17 is expressed in an opposite manner to Tsp: the sRNA is expressed in E phase while the protein is expressed only in PE phase (Figure 4). However, *tsp* seems to be transcribed in both E and PE phase (Figure 3C) and the relatively small increase in expression of the transcript between E and PE phase is insufficient to explain this observation. This suggest that Lpr17 blocks expression of Tsp. Since Tsp is important for survival at 55 °C, we hypothesize that Lpr17 should be repressed in this condition as well. Therefore, the expression of Lpr17 during thermal stress was analyzed by northern blot (Figure 5). The strains were grown to E phase in order to induced expression of Lpr17, and then subjected to a thermal stress at 55 °C. As seen in figure 5, Lpr17 sRNA was strongly repressed after 15 minutes at 55 °C in the WT and complemented strains. Lpr17 is repressed whenever Tsp is produced or needed and support the hypothesis that Lpr17 blocks translation of Tsp. Further experiments will be needed to confirm this possibility.

**Figure 5:**
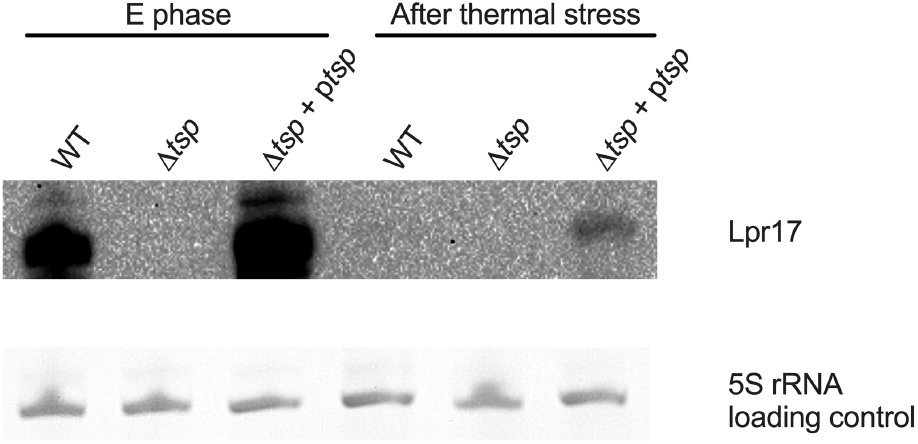
Lpr17 is repressed following thermal stress. The WT, *tsp* mutant (Δ*tsp*), and complemented strain (Δ*tsp* + p*tsp*) were grown to E phase in AYE broth and then subjected to heat shock at 55 °C for 15 minutes. Samples for RNA extraction were taken before (E phase) and after thermal stress for northern blot analysis.

### CpxR regulates *tsp* independently of Lpr17

A previous transcriptomic study has identified CpxR as a potential regulator of *tsp* in *Lp* (41) since the *tsp* gene was downregulated in the Δ*cpxR* mutant in the PE phase (41). In order to confirm that the downregulation observed also affects Tsp protein levels, the p*tsp*-his plasmid was electroporated in the Δ*cpxR* mutant and the expression of Tsp in the Δ*cpxR* mutant was analyzed by western blot (Figure 6A). In PE phase, Tsp is not expressed in the absence of CpxR. Since the Lpr17 sRNA seems to negatively regulates the expression of Tsp, we hypothesize that the deletion of CpxR may cause overexpression of Lpr17, which would inhibit Tsp expression. The expression of Lpr17 was therefore analyzed by northern blot (Figure 6B). The expression level of Lpr17 in PE phase in the Δ*cpxR* mutant is similar to the WT, indicating that the regulation of Tsp by CpxR is Lpr17-independent.

**Figure 6:**
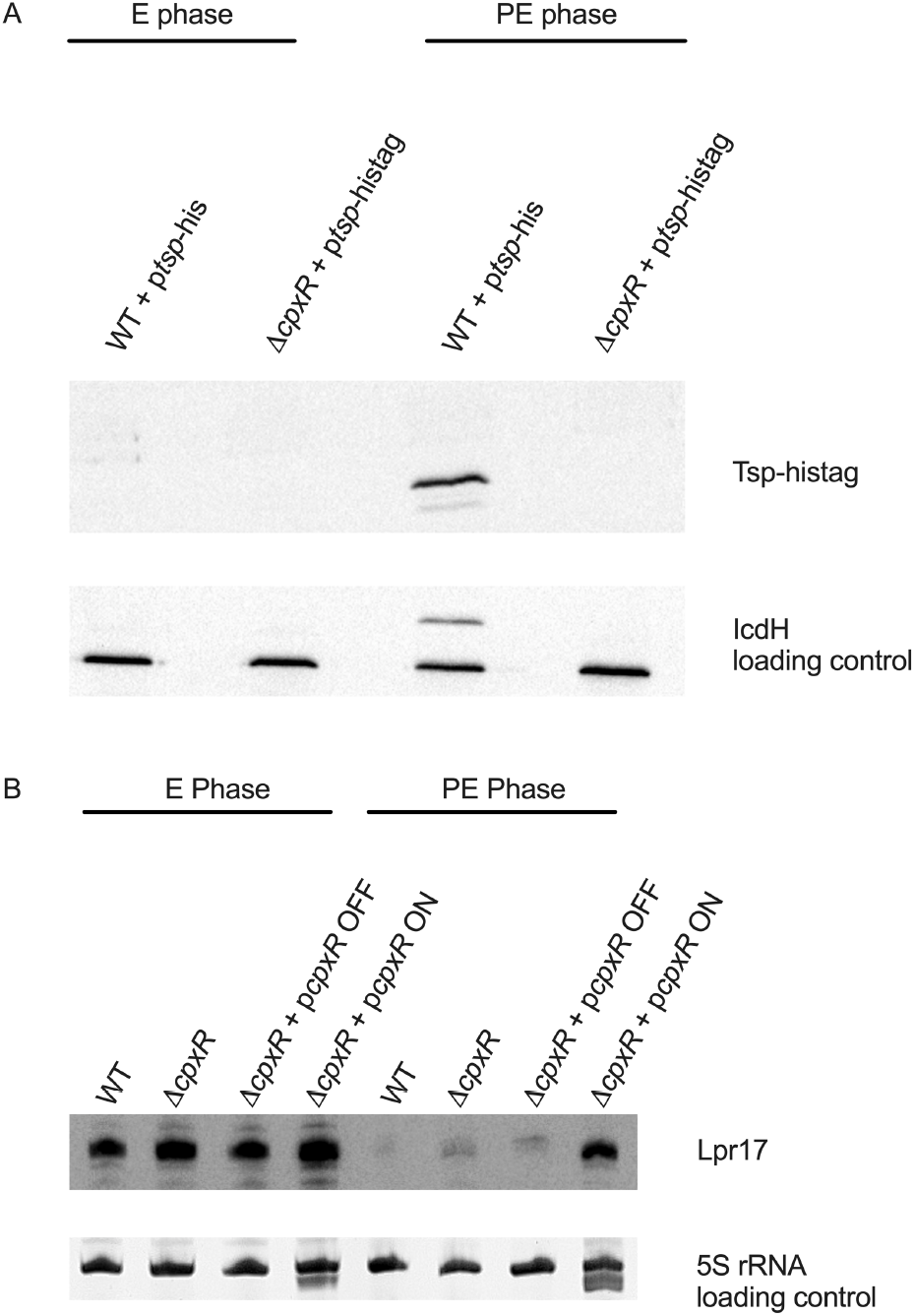
CpxR regulates Tsp in a Lpr17-independent manner. The WT and *cpxR* mutant Δ*cpxR* containing *ptsp-his* were grown to E and PE phase and analysed by (A) western blot and (B) northern blot. 5S rRNA and IcdH were used as a loading control for norther blot and western blot, respectively.

### The response regulator CpxR is important for surviving thermal stress

Since the Δ*cpxR* mutant produce no Tsp in PE phase, the ability of the *cpxR* mutant to survive thermal stress was investigated (Figure 7). The *cpxR* mutation was complemented in trans by cloning the Δ*cpxR* gene under the control of a *Ptac* promoter in the vector pMMB207c, since Δ*cpxR* is the fourth gene in a polycistronic mRNA. The *cpxR* mutant is unable to survive thermal stress compared to the WT and shows a much faster death than the wild-type. Basal expression from the vector without addition of IPTG was enough to complement the mutation (Δ*cpxR* + *pcpxR*). To ensure the complementation of the Δ*cpxR* mutant was due to the *cpxR* gene cloned, the *cpxR* mutant with the empty vector pMMB207c (Δ*cpxR* + pMMB207c) was also tested and showed a similar survival defect than the Δ*cpxR* mutant. The results were only significantly different after 15 minutes (*P* = 0.001). The CFU counts for the three replicates at the later timepoints showed large variation and the difference was not considered significantly different.

**Figure 7:**
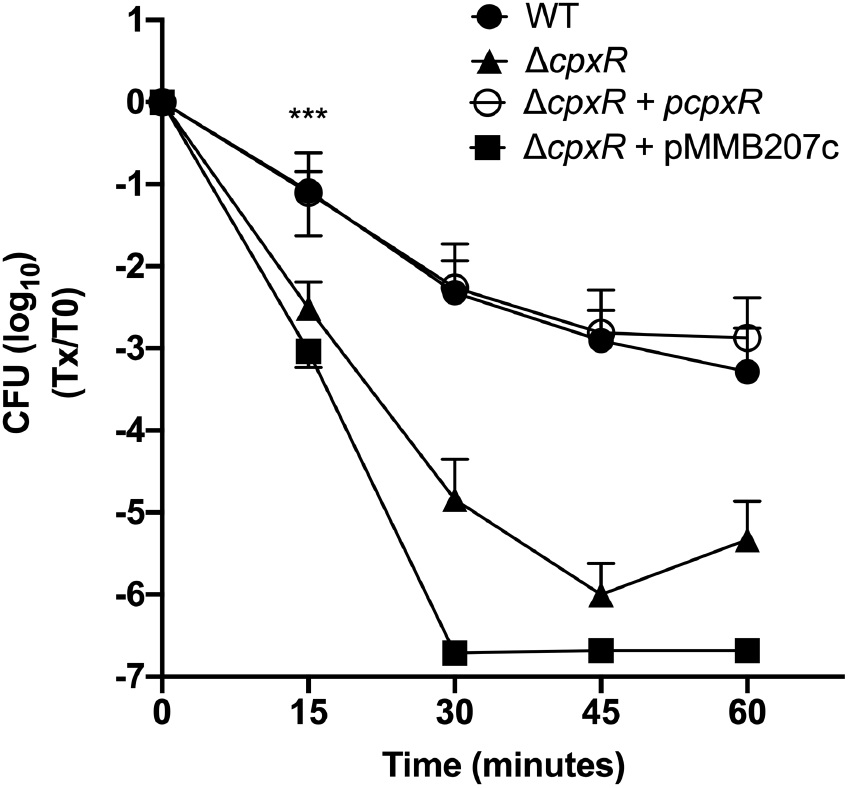
CpxR is required for *Lp* to survive a thermal stress. The WT, *cpxR* mutant (Δ*cpxR*), the *cpxR* mutant with the empty vector (Δ*cpxR* + pMMB207c), and the complemented strain (Δ*cpxR* + *pcpxR*) were suspended in Fraquil for 24 hours and then subjected to thermal stress at 55 °C. The survival of the strains was measured by CFU counts every 15 minutes from 0 to 60 minutes. The data shown represent the average of three independent replicate with standard deviation. A two-way ANOVA with a Tukey correction for multiple comparison was used to determine if the results were significantly different. The statistical difference shown in Figure 7 compares the WT and *tsp* mutant: ***, *P* < 0.005

## DISCUSSION

Similar to *tsp* mutants in other species, *Lp*’s *tsp* mutant is sensitive to thermal stress. The Δ*tsp* mutant is unable to tolerate 55 °C for 15 minutes (Figure 1). The *tsp* mutant is also sensitive to changes in temperatures. During infection of *V. vermiformis*, shifting the temperature from 25 °C to 37 °C impacts the ability of the *tsp* mutant to replicate (Figure 2B). The *tsp* mutant seems to suffer a delay in growth as well as the inability to reach the same level of growth achieved at 37 °C without a temperature shift (Figure 2C). The *tsp* mutant replicated to the same level as the WT if the temperature is maintained at 37 °C throughout the infection (Figure 2C). *Lp*’s ability to replicate within *V. vermiformis* despite shifts in temperature is likely important for its persistence in hot water distribution systems. Sporadic use of a system is likely to generate section of lower temperature, such as in dead ends, that momentarily gets warmer when the hot water flows through the system. In addition, increasing the temperature of the system is often used to disinfect hot water distribution systems (8). The amoeba cysts are able to survive treatments up to 80 °C, and *Lp*’s ability to survive and replicate efficiently once the temperature decreases is important for recolonization of the water system (3).

Possibly, the absence of Tsp causes a disturbance in the cell membrane resulting in increased sensitivity to thermal stress. This reasoning was previously suggested to explain decrease resistance to thermal stress of *tsp* mutants in *Borrelia burgdorferi, Escherichia coli*, and *Staphylococcus aureus* (20, 25, 50). Alternatively, accumulation of misfolded proteins in the *tsp* mutant could explain the inability to cope with thermal stress. It has previously been shown in other bacteria that Tsp degrades misfolded proteins that accumulate in the periplasm (26, 31, 32). Degradation of misfolded proteins increases the pool of amino acids available for *de novo* protein synthesis. Possibly, the accumulated misfolded proteins are not degraded in the absence of Tsp, leading to a decrease of the pool of amino acids (13, 14). This might explain the inability of the *tsp* mutant to grow as well as the WT following the temperature shift.

Despite not replicating as much as when the infection is carried out at constant temperature, the *tsp* mutant is still able to replicate within amoeba to some extent. This suggest that in the absence of Tsp, other proteases are able to degrade misfolded proteins, thereby relieving the stress. The expression of alternate proteases could explain the delay in growth compared to the WT observed following the temperature shift. In *E. coli*, the DegP periplasmic protease is important for surviving elevated temperatures and is responsible for degrading misfolded proteins and aggregated proteins in the periplasm (51–55). DegQ is a homolog of DegP that is also found in the periplasm and is responsible for degrading denatured proteins (56–58). In *Legionella*, the DegP homologue is required for surviving thermal stress, but is not required for infection of amoeba (59). The infection was done at a constant temperature, and therefore it is unknown if DegP plays a role during amoeba infection if a temperature change occurs (59). *Legionella* codes for a DegQ homologue, however its role in surviving thermal stress have not been investigated (60, 61). In the absence of *tsp*, it is possible that these proteases also contribute to removing the aggregated and misfolded proteins during the temperature that occurs during amoeba infection. However, subjecting the strain to 55 °C might be too big of a stress for the other proteases to compensate the lack of *tsp*. The presence of a N-terminal secretion signal in Tsp was found using two bioinformatic software, signalP and Phobius (62–64). This is consistent with *tsp*’s from other species that also have a N-terminal secretion signal required for transport to the periplasm (19). This suggests that *Lp*’s Tsp is located in the periplasm and could therefore degrade misfolded and aggregated proteins in the periplasm.

*P. aeruginosa’s* CtpA cleaves MepM, belonging to the M23 peptidase family. It is interesting to note that *tsp* is encoded downstream of *lpg0500*, which codes for a M23/M37 family peptidase. *Lp* codes for two other peptidases belonging to the M23/M37 family, *lpg0567* and *lpg0825*, the former being a homologue of *P. aeruginosa’*s MepM. In addition, *Lp* codes for at least 10 TPR repeat proteins, the same type of protein as the CtpA partner LcbA, suggesting one or some of these TPR proteins might be required for *Lp*’s Tsp’s activity, similarly to what has been reported in *P. aeruginosa* (65).

*Cis*-encoded sRNAs, such as Lpr17, typically regulate genes coded on the complementary strand (47). The expression pattern of Lpr17 and Tsp are opposite to each other, where the sRNA is only expressed in E phase (Figure 3B) while the Tsp protein is only expressed in PE phase (Figure 4B). This pattern of expression suggests a negative regulation of Tsp by Lpr17. This is further supported by the complete repression of Lpr17 during thermal stress (Figure 5). Furthermore, RT-PCR on cDNA extracted from E and PE phase (Figure 3C) shows the presence of the *tsp* mRNA transcript in both phases, whereas the protein is only detected in PE phase. Taken together, our results suggest that the Lpr17 sRNA inhibits translation of the protein, possibly by blocking the binding of ribosome to the ribosome binding site (RBS), without initiating degradation of the transcript (Figure 3A). This is reminiscent of several cis-encoded sRNA that overlap with the 5’ untranslated region of their target gene (66). Determining the exact mechanism would require additional experiments that are beyond the scope of the present study.

In *E. coli*, the CpxR/A two-component system regulates genes involved in dealing with envelope stress and misfolded proteins in the periplasm (67). In *E. coli*, CpxR upregulates the DegP periplasmic protease (68–70). In *Lp*, CpxR does not regulate DegP and DegQ in E and PE phase, but seems essential for expression of Tsp (41). Therefore, the inability of the *cpxR* mutant to tolerate thermal stress could rely on the absence of Tsp; however, complementation of *cpxR* defect by expressing *tsp* in *trans* was unsuccessful, suggesting that CpxR regulates other determinants of thermal stress resistance. It is unclear if CpxR directly binds to the promoter region or if the regulation of *tsp* is indirect. Additional experiments are required to investigate the exact mechanism.

In conclusion, we have demonstrated that Tsp is necessary for survival of thermal stress in water and during intracellular growth. Such situations are likely to be encountered by *Lp* in its normal environment, which makes Tsp a critical genetic determinant for survival and growth in water systems. Our results show that *tsp* is likely regulated by a complex network consisting minimally of Lpr17 and CpxR. Finally, we have determined that CpxR is an important regulator of thermal stress tolerance in *Lp*.

## MATERIAL AND METHODS

### Bacterial strains and media

Table 1 describes the bacterial strains used in this study. The WT *Lp* strain used in this study is KS79, a Δ*comR* mutant of the JR32 strain, which allows the strain to be constitutively competent (71). JR32 is a Philadelphia-1 derivative that is salt-sensitive, streptomycin-resistant, and restriction negative (72). *Lp* strains were cultured on CYE agar (ACES-buffered charcoal yeast extract) supplemented with 0.25 mg ml^−1^ of L-cysteine and 0.4 mg ml^−1^ of ferric pyrophosphate (73). Strains were cultured in AYE broth, which is CYE lacking charcoal and agar. If required, media were supplemented with 25 μg ml^−1^ of kanamycin and 5 μg ml^−1^ of chloramphenicol (73).

**Table 1:**
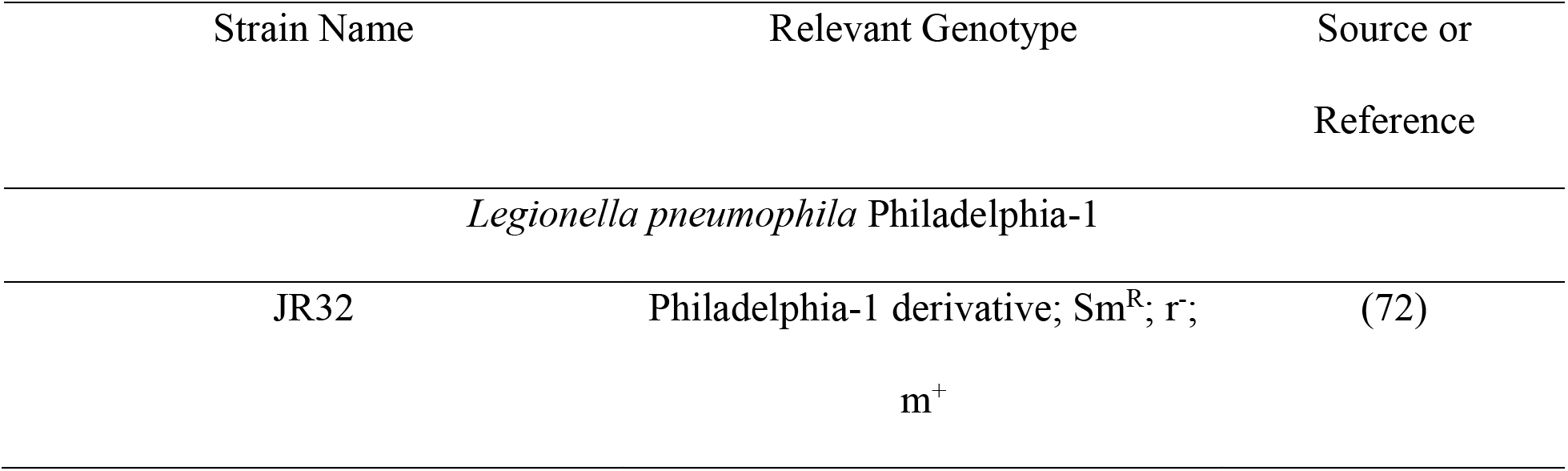
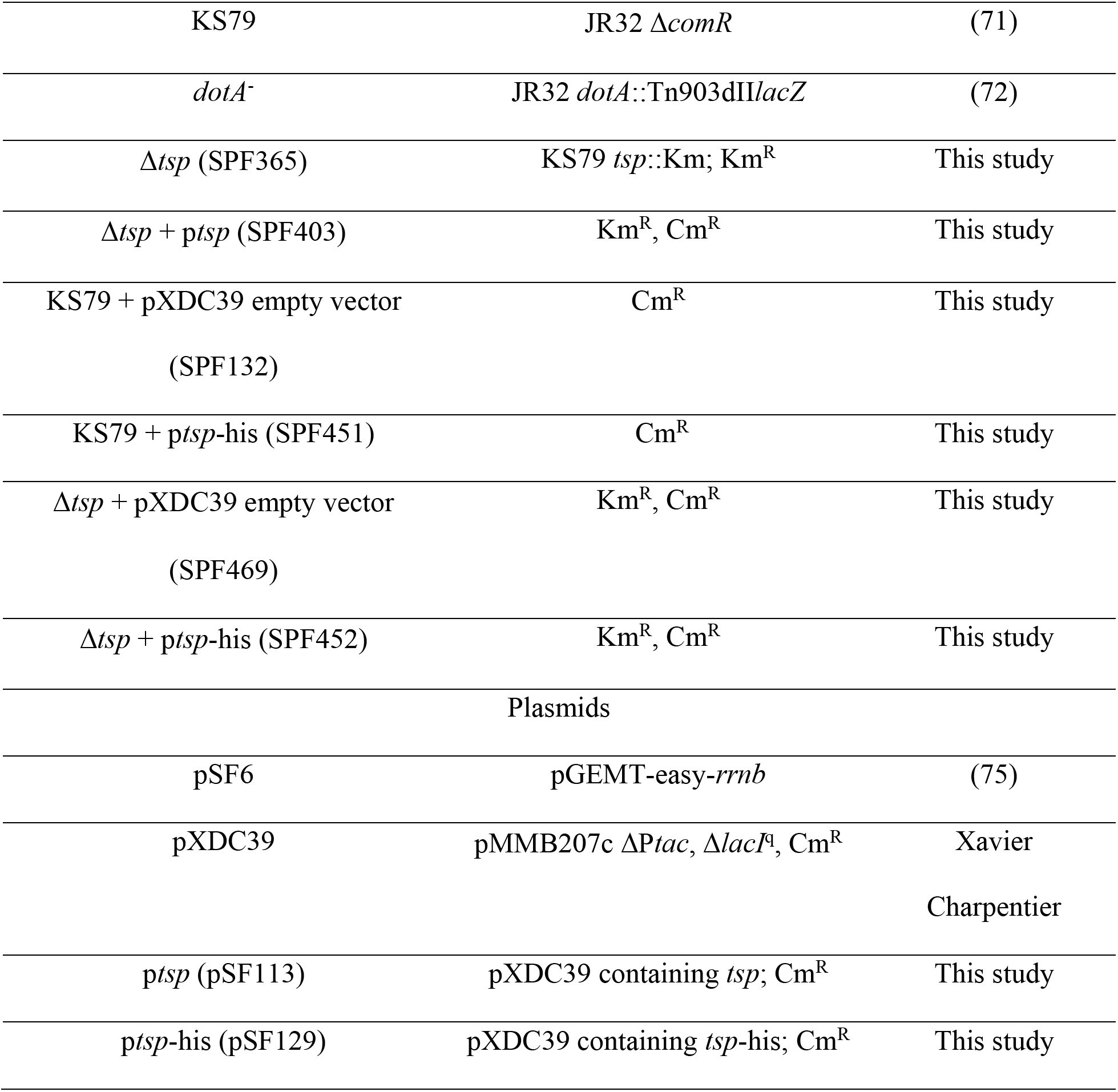
Strains used in this study

### Deletion of *tsp* and complementation of the mutant

The *tsp* gene was replaced with a kanamycin resistance cassette by allelic exchange, as previously described, to construct the deletion mutant strain (74). PCR primers are described in Table 2. A 1-kb fragment upstream of *tsp* was amplified using primers Lpg499-UF and Lpg499-UR. A 1-kb fragment downstream of *tsp* was amplified using primers Lpg499-DF and Lpg499-DR. The kanamycin cassette was amplified from the pSF6 plasmid (75) using primers Lpg499-Km-F and Lpg499-Km-R. Each 1-kb fragment was purified using a gel extraction kit (Qiagen) and were ligated by PCR using primers Lpg499-DR and Lpg499-UF to generate a 3-kb fragment that was purified by a gel extraction kit (Qiagen). The purified 3-kb fragment was introduced into the KS79 strain by natural transformation (76) and the recombinants were selected on CYE agar supplemented with kanamycin. PCR was used to confirm the allelic exchange and the Δ*tsp* mutant strain was named SPF365.

**Table 2:**
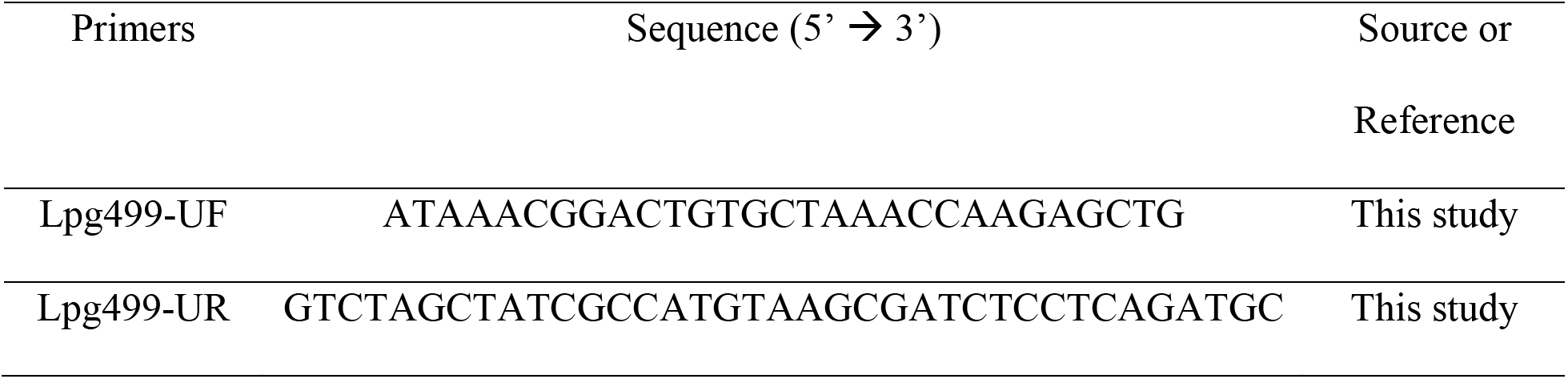
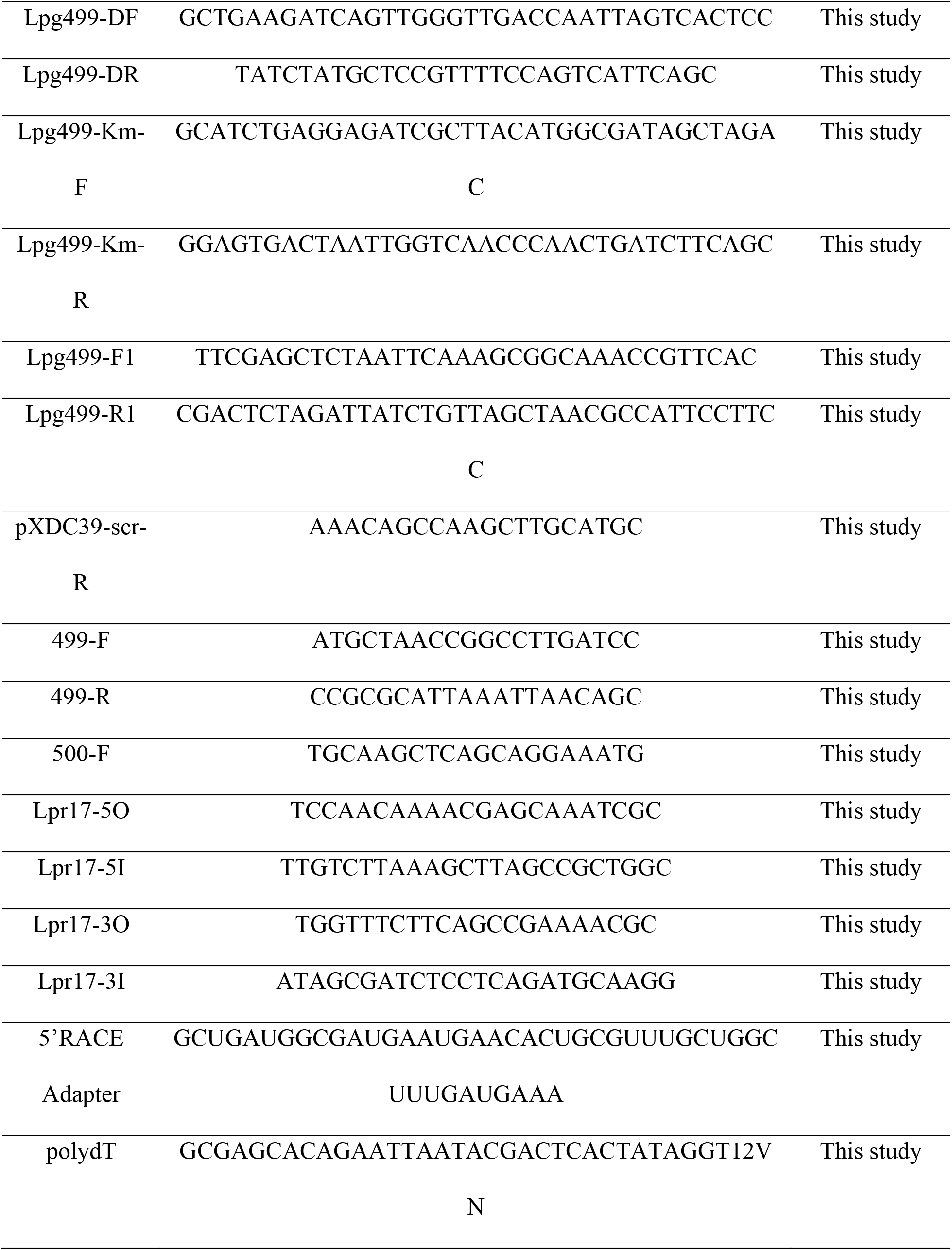
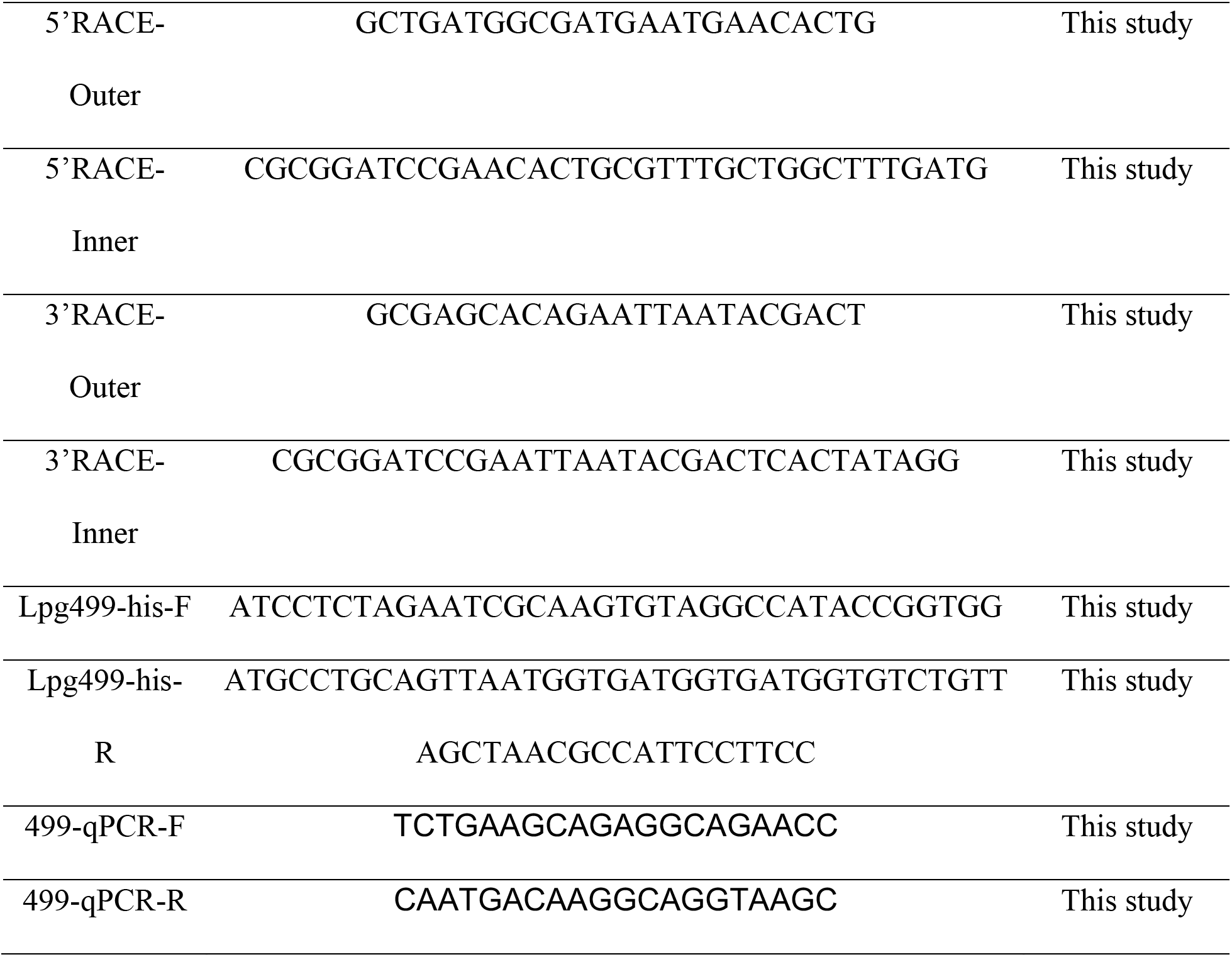
Primers used in this study

The *tsp* mutation was complemented in trans by amplifying the *tsp* gene with its native promoter from the KS79 WT genomic DNA using primers Lpg499-F1 and Lpg499-R1. The Lpg499-F1 primer is located 441 nucleotides upstream of *tsp*’s ATG and the amplicon includes both the *tsp* and *lpr17* genes. The amplicon was cloned in the pXDC39 plasmid vector using the restriction enzymes SacI (New England Biolabs) and XbaI (New England Biolabs). The restriction digestion was done at 37 °C for 1 hour, column purified (Qiagen), and ligated for 2 hours at room temperature using T4 DNA ligase (New England Biolabs). The ligation product was transformed into *E. coli* DH5α and the transformants were selected on LB agar containing 25 μg/ml of chloramphenicol. Cloning was confirmed by PCR using primers Lpg499-F1 and pXDC39-scr-R. The recombinant plasmid (p*tsp*) was extracted and electroporated into the Δ*tsp* mutant strain. The transformants were selected on CYE agar containing 5 μg ml^−1^ of chloramphenicol, then patched on CYE agar containing 25 μg ml^−1^ of kanamycin and 5 μg ml^−1^ of chloramphenicol. Screening was done by PCR using primers Lpg499-F1 and pXDC39-scr-R and the resulting strain was called SPF403.

### Thermal stress

*Lp* strains were grown on CYE agar for 3 days at 37 °C. A suspension was prepared from a few colonies in Fraquil, an artificial freshwater medium (77, 78) at an OD_600_ of 1.0. The suspension was washed three time with Fraquil to remove any trace of nutrients. The strains were incubated at 25 °C for 24 hours in 25 cm^2^ cell culture flasks (Sarstedt). The strains were then diluted 1:10 in Fraquil and 1 ml were distributed in 13-ml tubes (Sarstedt) and incubated in a 55 °C water bath. Individual tubes were used for each time point. Tubes were removed from the water bath for CFU counts on CYE agar to determine the survival of the strains. CFU counts were done at 0, 15, 30, 45, and 60 minutes.

### *Vermamoeba vermiformis* culture and infection

*V. vermiformis* cells were grown at room temperature in 75 cm^2^ cell culture flasks (Sarstedt) in modified PYNFH medium (ATCC medium 1034) and passaged when confluence was reached. The amoebas were centrifuged at 200 g for 10 minutes, supernatant discarded, and cells were resuspended in fresh modified PYNFH medium (ATCC medium 1034) and incubated for three days prior to infection. On the day the infection was to be carried, the cells were centrifuged at 200 g for 10 minutes and resuspended at a concentration of 5.0 × 10^5^ cells/ml in modified PYNFH lacking FBS and the buffer solution. *Legionella* is able to grow in modified PYNFH containing FBS and the buffer solution, therefore they were omitted during the infection to ensure that the growth observed is due to intracellular multiplication. 1 ml of cells were seeded in each well of the 24-well plate (Sarstedt).

Bacterial strains were grown on CYE agar with antibiotics for 3 days at 37 °C. On the day of the infection, the bacteria were suspended in Fraquil at an OD_600_ of 0.1 and then diluted 1:10 in Fraquil. 5 μl of the 1:10 dilution was added to each well in triplicate in order to have a starting MOI of 0.1. The infection was carried at room temperature for 5 days, at 37 °C for 5 days, and at 25 °C for 2 days and then at 37 °C for an additional 3 days. CFU counts were performed on a daily basis.

### RNA extraction

TRIzol (ThermoFisher Scientific) was used for RNA extraction according to the manufacturer’s protocol. Briefly, 200 μl of chloroform (ThermoFisher Scientific) was added to the bacterial pellet suspended in TRIzol, shaken, and incubated for 3 minutes at room temperature. The sample was added to a Phase-lock Gel Heavy 2 ml tube (VWR) and centrifuged at 12,000 × *g* for 15 minutes at room temperature. Approximately 600 μl of the supernatant was transferred to a microtube containing 600 μl of 100% isopropanol (ThermoFisher Scientific) and 5 μg of glycogen (ThermoFisher Scientific). Following incubation at room temperature for 10 minutes, the tubes were centrifuged at 17,000 × g for 10 minutes at 4 °C. The supernatant was removed and 1 ml of 75 % ice-cold ethanol (Greenfield Global) was added to the RNA pellet. The tubes were centrifuged at 17,000 × g for 10 minutes at 4 °C, the supernatant removed, the pellet air dried and resuspended in nuclease-free water (ThermoFisher Scientific). The RNA was quantified using a UV spectrophotometer (ThermoFisher Scientific).

### Northern Blotting

The expression of the Lpr17 sRNA was examined by northern blot as previously described (15). The strains were grown on CYE agar with required antibiotics for 3 days at 37 °C, inoculated in AYE broth with required antibiotics and grown at 37 °C with shaking at 250 rpm. Aliquots of 10 ml of exponential phase culture (OD_600_ 0.5-1.0) and 5 ml of PE phase culture (OD_600_ >3) were centrifuged at 4,000 g for 10 minutes, the supernatant removed, and the bacterial pellet resuspended in 1 ml of TRIzol reagent (ThermoFisher Scientific). RNA extraction was carried out as mentioned above. 5 μg of RNA was loaded on a 6 % Tris-borate-EDTA-urea polyacrylamide gel and the samples were migrated at 180 mV. The RNA was transferred to a positively charged nylon membrane (ThermoFisher Scientific) using a semidry gel blotting system (Biorad) for 20 minutes at 200 mA. The membrane was prehybridized in ULTRAhyb-Oligo Hybridization buffer (ThermoFisher Scientific) for 1 hour at 37 °C in a rotating chamber. 5 nM of 5’ biotinylated Lpr10 probe (Integrated DNA Technologies) was added to the prehybridization buffer and the membrane was incubated overnight at 37 °C in the rotating chamber. The membrane was twice washed with 2X SSC (0.15 M NaCl and 0.015 M sodium citrate) and 0.5 % SDS (Bio-Rad) for 30 minutes at 37 °C. The probes were detected with the Chemiluminescent Nucleic Acid Detection Module (ThermoFisher Scientific).

### RT-PCR

RNA extracted from the KS79 WT strain grown to E and PE phase was treated with DNase (ThermoFisher Scientific). 1 μg of DNase treated RNA was reverse transcribed with Protoscript II (New England Biolabs) and a no reverse transcriptase (no RT) was included by replacing the reverse transcriptase by nuclease-free water. The PCR was performed on cDNA, no RT control, WT gDNA, and a no template control (water) using primers described in Table 2. The amplicon was analysed on a 1 % agarose gel.

### Quantitative PCR

RNA was extracted and cDNA synthesized as described above. Following cDNA synthesis, qPCR was done with primers 499-qPCR-F and 499-qPCR-R. qPCR was performed using iTaq Universal SYBR Green supermix (Bio Rad) according to the manufacturer’s protocol. The efficiency of the primer pairs was determined using dilution series of gDNA. Ct values were normalized to the 16S rRNA as described previously (79).

### Cloning of Tsp with polyhistidine tag

The polyhistidine tag was added to *tsp* by PCR. *tsp* along with its native promoter was amplified using primers Lpg499-his-F and Lpg499-his-R, the latter containing a polyhistidine tag. The amplicon was cloned in the pXDC39 plasmid vector using XbaI (New England Biolabs) and PstI (New England Biolabs). The restriction digestion was carried at 37 °C for 1 hour, followed by column purification (Qiagen), and ligation at room temperature for 2 hours using T4 DNA ligase (New England Biolabs). The ligation product was transformed into *E. coli* DH5-a and the transformants were selected on LB agar supplemented with 25 μg ml^−1^ of chloramphenicol. The cloning was confirmed by PCR using primers Lpg499-his-F and pXDC39-scr-R. The plasmid (p*tsp*-his) was extracted and electroporated into KS79 and the *tsp* mutant and transformants were selected on CYE containing 5 μg ml^−1^ of chloramphenicol. The KS79 p*tsp*-his transformants were patched on CYE containing 5 μg ml^−1^ of chloramphenicol while the *tsp* mutant p*tsp*-his transformants were patched on CYE containing 25 μg ml^−1^ of kanamycin and 5 μg ml^−1^ of chloramphenicol. Screening was done by PCR using primers Lpg499-his-F and pXDC39-scr-R. The pXDC39 empty vector was also electroporated in the KS79 WT and the *tsp* mutant and serve as controls.

### Western blot

Strains were grown to E and PE phase in AYE broth. Aliquots of 1.5 ml were centrifuged at 17,000 g for 3 minutes, the supernatant was decanted, and the cell pellet resuspended in 200 μl of 1X sample buffer (10 % glycerol, 62.5 mM Tris-HCl pH 6.8, 2.5 % SDS, 0.002 % bromophenol blue, 5 % β-mercaptoethanol). The samples were boiled for 5 minutes and sonicated for 15 minutes in an ice-cold water bath using Ultrasonic Cleaner (Cole-Palmer). Samples were then centrifuged for 15 minutes at 17,000 × *g* at 4 °C. The supernatant was transferred to a new tube and stored at −20 °C until it was ready to use. The protein samples were standardized by adjusting the final OD_600_ to 0.5 by diluting the samples with 1X sample buffer. 15 μl of the standardized protein sample was loaded on a 12.5 % polyacrylamide gel and samples were migrated at 100 V. The proteins were transferred onto a PVDF membrane (Bio Rad) at 16 V for 24 hours. The membrane was blocked for 30 minutes with a 5 % milk protein solution. Tsp-his was detected with the Anti-6X His tag antibody (ThermoFisher Scientific), which is already conjugated with HRP and does not require a secondary antibody for detection. The blot was incubated with this antibody at room temperature for 2 hours followed. The IcdH antibody (Sigma) was used as a loading control. In this case the blot was incubated with the IcdH primary antibody at room temperature for 2 hours and with the secondary antibody (anti-rabbit HRP, Sigma) was done at room temperature for 30 minutes. Antibodies were detected using ECL Prime Western Blotting Detection Reagents (GE Healthcare).

## ACKNOWLEDGMENTS

The pXDC39 plasmid is a kind gift from Dr. Xavier Charpentier. This study was supported by CIHR Open Operating Grant #142208 to SPF. JS was supported by a FRQNT Doctoral scholarship and a CRIPA scholarship supported by the Fonds de recherche du Québec - Nature et technologies n°RS-170946.

